# A dual involvement of Protocadherin-18a in stromal cell development guides the formation of a functional hematopoietic niche

**DOI:** 10.1101/2021.10.19.465038

**Authors:** Anne-Lou Touret, Catherine Vivier, Anne Schmidt, Philippe Herbomel, Emi Murayama

## Abstract

Hematopoietic stem and progenitor cells (HSPCs) emerge from the aorta and migrate to the caudal hematopoietic tissue (CHT) of zebrafish larvae, the hematopoietic equivalent of the mammalian fetal liver, for their proliferation and differentiation. We previously reported that somite-derived stromal cells were a key component of the CHT niche. Here we found that the cell adhesion protein protocadherin-18a (Pcdh18a) is expressed in the stromal cell progenitors (SCPs) emigrating from somites toward the future CHT. Deletion of most of the intracellular domain of Pcdh18a caused a decrease in the number of SCPs, the directionality of their migration, and the cell-contact mediated repulsion that normally occurs between migrating SCPs. These defects were followed by abnormal morphogenesis of the venous plexus that forms the CHT framework, and the inability of the resulting CHT to function as a niche for HSPCs. Finally, we found that the extracellular domain of Pcdh18a mediates trans heterophilic adhesion of stromal cells to endothelial cells *in vivo* and thereby the reticular vs. perivascular fate of SCPs. Our study demonstrates that Pcdh18a expression in SCPs is essential for the proper development of the hematopoietic niche.

## Introduction

In vertebrates, hematopoiesis occurs throughout life from hematopoietic stem and progenitor cells (HSPCs) in dedicated niches, such as the bone marrow in adult tetrapods. Such niches are typically structured by two kind of cells, vascular endothelial cells and mesenchymal stromal cells, among which several sub-types have been discerned (reviewed in Gao et al., 2018). Developmentally, the definitive HSCs arise from the embryo’s aorta, and then migrate to their first niche, the fetal liver in mammals, in which they undergo expansion and differentiation into all hematopoietic cell types. Then they migrate to the bone marrow to establish lifelong hematopoiesis. While adult-type hematopoiesis in fish takes place in the kidney marrow, we found that in fish larvae, the aorta-derived definitive HSPCs first home to a dedicated niche in the tail, which we named the caudal hematopoietic tissue (CHT), that acts as the hematopoietic homologue of the fetal liver in mammals (Murayama et al., 2006). We found that the CHT niche consists in two main components, blood vessels - namely the caudal venous plexus (CVP) - and stromal cells, that interconnect the vascular segments of the CVP. More recently, we found that these stromal cells arise from cell clusters formed within the ventral side of the caudal somites, that undergo an epithelial mesenchymal transition (EMT) and migrate ventralwards into the presumptive CHT (Murayama et al., 2015). When the maturation of these stromal cell progenitors (SCPs) is impaired, the formation of the CHT niche is affected, resulting in failure to maintain HSPCs and support definitive hematopoiesis (Murayama et al., 2015). At the same time and in the same area, endothelial cells also migrate ventralwards from the primordial caudal vein to form the caudal venous plexus through sprouting angiogenesis (Wakayama et al., 2015). Previously, we reported that the migration of SCPs and vascular endothelial cells during CVP formation seemed to be synchronized, but the molecular mechanisms were not known (Murayama et al., 2015).

Protocadherins (Pcdhs) are cell adhesion molecules that comprise the largest family of proteins within the cadherin superfamily and are subdivided into clustered and non-clustered Pcdhs. The non-clustered Pcdhs are evolutionarily conserved and vary in their regional pattern of expression, prominently within the nervous system (Vanhalst et al., 2005). They further divide into δ1 and δ2 subfamilies based on differences in the number of extracellular cadherin (EC) repeats as well as in the intracellular domain (ICD) (Redies et al., 2005). 2-Pcdhs comprise Pcdh8, Pcdh10, Pcdh17, Pcdh18 and Pcdh19, they have six EC repeats, and their ICD bears two conserved motifs (CM1 and CM2) (Vanhalst et al., 2005). δ-Pcdhs seem to mediate homophilic cell-cell interactions, although binding appears to be weaker than that of classical cadherins (Harrison et al., 2020; Tai et al., 2010). In addition, δ 2-Pcdhs were shown to mediate heterophilic *cis* interactions with classical cadherins (Biswas et al., 2010; Chen et al., 2009; Kraft et al., 2012). It still not established whether they can engage heterophilic *trans* interactions. Pcdh17 and Pcdh19 carry a conserved RGD motif in their extracellular domain (EC2), suggesting that they may interact with integrins (Jontes, 2016), but an actual molecular interaction has yet to be demonstrated. Some Pcdhs have been reported to be involved in the fine tuning of adhesion versus repulsion. For example, a deletion of the ICD of Pcdh17 led to a defect in cellular repulsion, resulting in axonal clumping of abducens motor neurons (Asakawa and Kawakami, 2018). The ICD of Pcdhs is known to activate downstream signaling cascades (Pancho et al., 2020). Several Pcdhs are involved in the regulation of actin cytoskeletal dynamics through the WAVE regulatory complex (WRC), and the WRC interacting receptor sequence (WIRS) is highly conserved among Pcdhs of the δ2 subfamily (Chen et al., 2014). Furthermore, Pcdh18 was identified as a binding partner for Disabled-1 (Dab1), a component of the Reelin signaling pathway that mediates neuronal migration during cortical lamination (Homayouni et al., 2001). Several studies reported that the downstream effectors of the non-clustered Pcdhs are involved in cell movements and motility (Nakao et al., 2008; Biswas et al., 2014; Hayashi et al., 2014).

Here, we report the identification of Pcdh18a as a molecule that is continuously expressed in the SCPs during their emergence and migration in zebrafish embryos and is essential for the formation of a functional hematopoietic niche. Pcdh18a affects the migration of SCPs, notably the repulsion among leader cells, the number of SCPs, and their interactions with endothelial cells, the other main component of the CHT niche. It mediates stromal cell interaction with the endothelium, and thereby conditions the subsequent differentiation fate of SCPs.

## Results

### Somite-derived stromal cell progenitors migrate alongside vascular endothelial cells

We previously reported that cell clusters containing stromal cell progenitors (SCPs) appeared at the ventral side of caudal somites from 21-22 hpf (Murayama et al., 2015). Somites are formed one by one every 30 minutes in rostro-caudal direction, and then progressively mature (Stickney et al., 2000). They initially display an epithelial structure on their entire surface; then cell clusters consisting of about 15 to 20 cells per somite become apparent on the ventral side, and from these clusters the SCPs soon emigrate as strings in ventral direction (Murayama et al., 2015). At about the same time, vascular endothelial cells sprout from the primordial caudal vein (PV) at the midline and migrate ventralwards together with the SCPs. To examine the spatiotemporal migration path of SCPs in the caudal region relative to endothelial cells, we firstly performed live imaging using Tg*(pax3a:GFP; kdrl:ras-mCherry)* or Tg*(ET37:GFP; kdrl:ras-mCherry)* double transgenic embryos. The pax3a:GFP and ET37:GFP transgenes are initially expressed weakly throughout the somites, then pax3a:GFP becomes more strongly expressed in the lateral and ventral cell layers (Lee et al., 2013) which comprise the SCP clusters; then both transgenes are well expressed in the emigrating SCPs. The kdrl:mCherry transgene specifically labels endothelial cells. Confocal time-lapse imaging of the caudal region of double transgenic embryos in lateral view followed by digital reconstruction of transverse sections revealed that by 23-25 hpf, SCPs that delaminated from the somites then migrating ventralwards on the outer surface of the PV (Fig. 1A-a,b; Movie 1). By 27 hpf, they migrated further ventrally together with endothelial cells extending from the ventral wall of the PV (Fig. 1B,a and Movie 1), which by 36 hpf had fully organized into a functional caudal venous plexus (CVP) including the definitive caudal vein (CV) as its ventral-most segment (Fig. 1A-c,d). Meanwhile, SCPs/SCs had settled in between the vascular segments of the CVP, and some of them had migrated further ventrally into the caudal fin to become fin mesenchymal cells (FMCs) (Lee et al., 2013) (Fig. 1A-c,d, purple arrows). *In vivo* imaging at higher magnification revealed that during CVP morphogenesis, migrating SCPs and vascular endothelial cells dynamically stretched out filopodia through which they interacted (Fig. 1B,b-d), then endothelial cells appeared to join and form new lumenized vascular segments around the neighboring SCPs (Movie 1).

**Figure 1.**
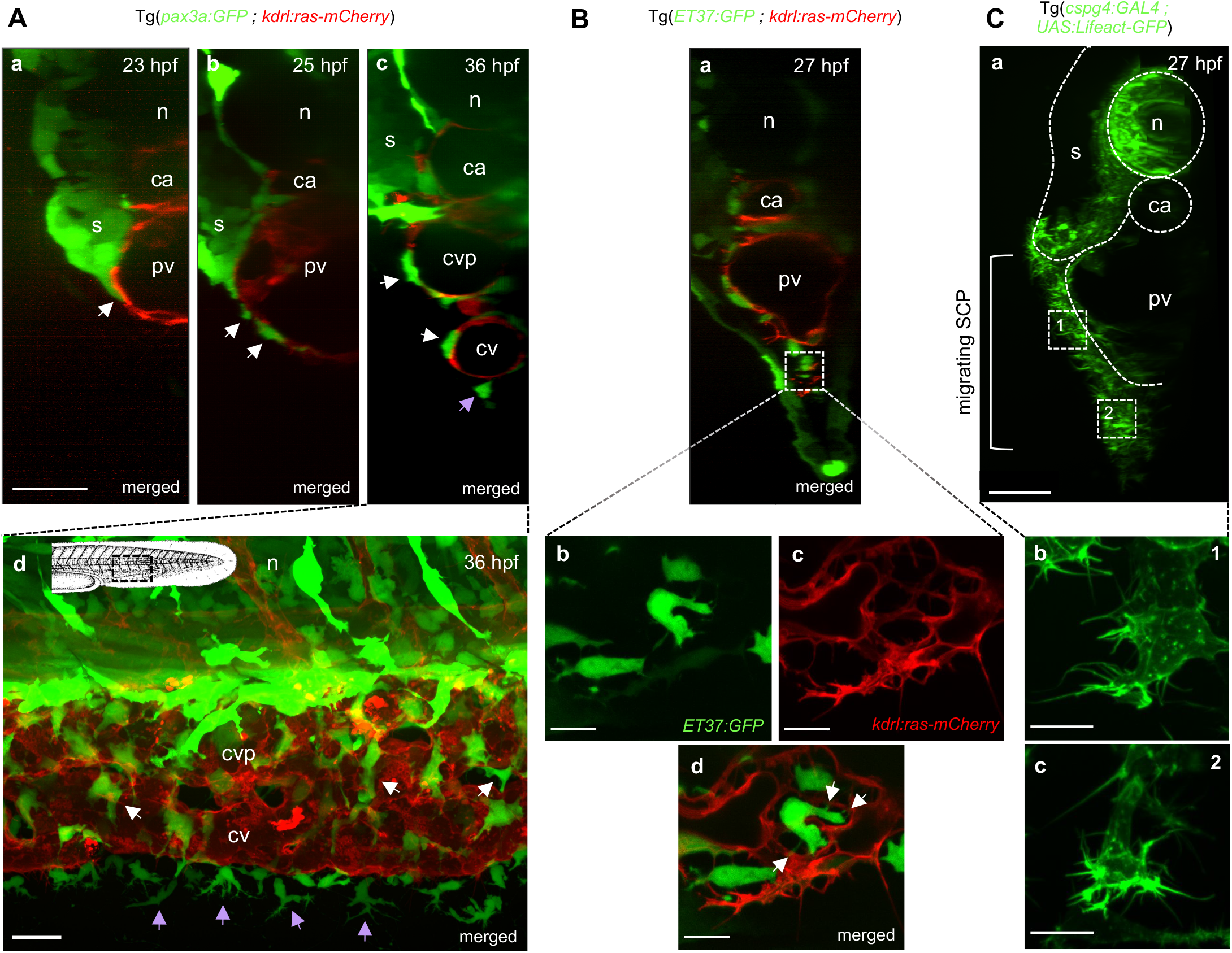
Stromal-endothelial cell interactions during SCP migration and caudal vein plexus formation. **(A)** Confocal spinning-disk acquisitions of live transgenic Tg(*pax3a:GFP; kdrl:ras-mCherry*) embryos. **(a**,**b**,**c)** Optical transverse sections reconstituted by Imaris software for wild-type embryos at 23, 25 and 36 hpf. White arrows indicate SCPs migrating on the primordial vein or within the forming caudal vein plexus; purple arrows indicate fin mesenchymal cells (FMCs). Scale bar, 30 µm. **(d)** Projection of the caudal region at 36 hpf. GFP^high^ and GFP^dim^ cells correspond to neural crest derived pigment cells and SCP derivatives, respectively. Scale bar, 20 µm. **(B)** Confocal spinning-disk images of live transgenic Tg(*ET37:GFP; kdrl:ras-mCherry*) embryos at 27 hpf. **(a)** Optical transverse section. **(b**,**c**,**d)** Magnified projections from 3 confocal planes spaced by 0.6 µm at the region indicated by a dotted square in (a), ventral to the primordial vein where sprouting angiogenesis is active to form the future caudal vein plexus. Active protrusions and filopodia are observed in migrating endothelial cells **(c)** and a stromal cell interacts with an endothelial cell through some filopodia (**d**, white arrows). Scale bars, 10 µm. **(C)** Confocal spinning-disk images of live Tg(*cspg4:GAL4; UAS:Lifeact-GFP*) embryo at 27 hpf. **(a)** Optical transverse section. SCPs first migrate in a 2D mode in close contact with the primordial vein (dotted square indicated by **1**) then in a more 3D mode when they co-migrate with developing endothelial cells (dotted square indicated by **2**). Scale bar, 20 µm. **(b, c)** Magnified images of SCPs in 2D **(b)** and 3D **(c)** migration modes. Scale bars, 10 µm. n, notochord; s, somites; ca, caudal artery; pv, primordial vein; cvp, caudal vein plexus; cv, caudal vein.

To further investigate the morphodynamics of migrating SCPs, we switched to a new transgenic driver line created by us (Murayama et al., manuscript in preparation), TgBAC(*cspg4:GAL4*), which drives specific expression of Gal4 in SCPs and stromal cells (SCs) in addition to the notochord, and we combined it with a Tg(*UAS:Lifeact-GFP*) reporter line where Lifeact-eGFP highlights F-actin dynamics *in vivo*. SCPs migrating on the outer surface of the PV and further ventrally extended filopodia toward their leading edge and in many other directions, and formed an extensive interconnected network (Fig. 1C, Movie 2).

### The cytoplasmic domain of Pcdh18a is involved in stromal cell progenitor migration

To understand the molecular mechanism of SCP emigration from the somites, we performed whole-mount in situ hybridization (WISH) analysis for genes encoding cell adhesion molecules, generally involved in cell migration, during SCP emergence in the caudal somites and the ensuing migration period. We found no cadherin expression specifically in SCPs, so we moved to the protocadherin family. We found that *pcdh18a*, encoding Protocadherin-18a, was expressed in developing SCPs (Fig. 2A). *pcdh18a* expression was first observed in the SCP clusters at the ventral side of caudal somites (Fig. S1a,a’), then also in the migrating SCPs derived from them (Fig. S1-b,b’), and so at least until 35 hpf (Fig. S1c,c’).

**Figure 2.**
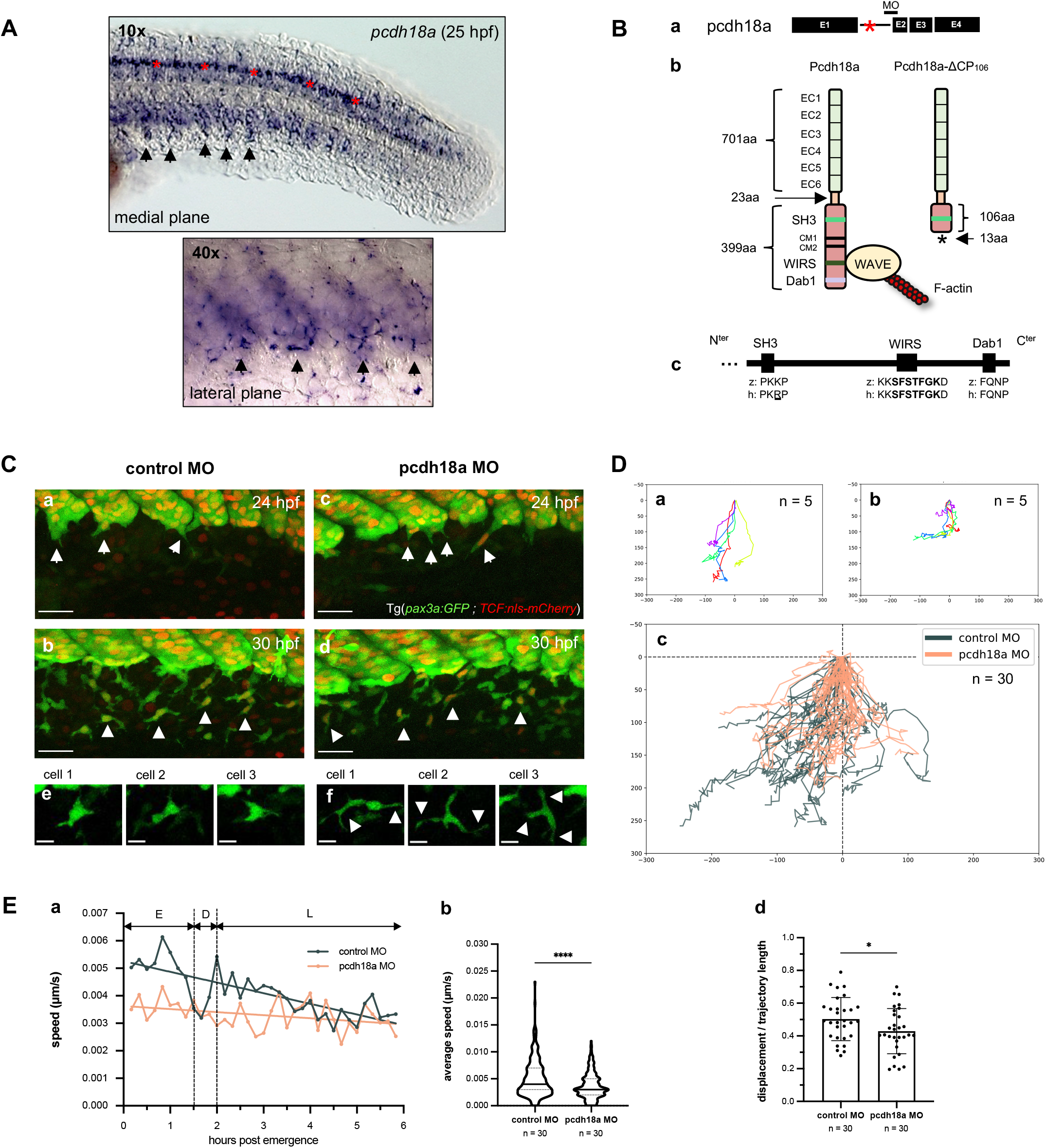
Pcdh18a is involved in the migration of SCPs. **(A)** VE-DIC microscopy of whole-mount in situ hybridization of *pcdh18a* at 25 hpf. The ventral side of caudal somites and SCPs migrating from the clusters (arrows) are labeled by pcdh18a mRNA probe from the 5^th^ somite counted from the tailbud and in the more rostral (more mature) somites. Some neurons in the spinal cord are also labelled (red asterisks). **(B-a)** The splice-blocking MO used in this study targets the intron1-exon2 junction of *pcdh18a*. **(B-b)** Injection of this MO leads to retention of the first intron and a premature STOP codon in it, hence to a protein truncated after the 106^th^ amino acid of the intracellular domain (ICD) of Pcdh18a. **(B-c)** Alignment of SH3, WIRS and Dab1 domains in the ICD of human PCDH18 and zebrafish Pcdh18a. **(C)** Time-lapse confocal imaging of Tg(*pax3a:GFP; TCF:nls-mCherry*) control or pcdh18a-ΔCP_106_ morphant embryos. Maximum projections were extracted at 24 **(a**,**c)** and 30 hpf **(b**,**d)**. Arrows and arrowheads indicate delaminating and migrating SCPs/SCs, respectively; scale bars, 30 µm. **(e**,**f)** Snapshots of single SCPs during mid-migration (26-29 hpf); scale bars, 10 µm. **(D-a**,**b)** Individual tracks of 5 leader SCPs obtained from the time-lapse imaging for a control embryo **(a)** and a pcdh18a-ΔCP_106_ morphant embryo **(b)**; cell positions were plotted every 10 minutes during 6 hours; in a trajectory plot, all (x,y) coordinates of the SCPs’ starting points (in the clusters) are set to (0,0) and negative or positive values on the X axis indicate anterior or posterior migration, respectively **(D-c)** Overlay of individual tracks of the 30 leader cells followed in control (black) and pcdh18a-ΔCP_106_ morphant (red) embryos. n_exp_=3, n_embryos_=6 for controls and morphants. **(D-d)** Trajectory straightness (ratio of the net displacement over trajectory length) of the tracked leader SCPs (mean±SD; ^*^, P = 0.0378; Student’s t test). **(E-a)** Average speed of the tracked leader SCPs over time. E, early migration; D, decision phase; L, late migration. **(E-b)** Violin plot of the speed of leader SCPs during phase E of their migration (median±SD; ^****^, P<0.0001; Mann-Whitney test).

To investigate the possible involvement of Pcdh18a in SCP emigration from the somites, we firstly used an antisense morpholino (MO). Previous studies found that morpholinos blocking Pcdh18a translation caused defects in the early development of zebrafish embryos and lethality by mid-somitogenesis (Aamar and Dawid, 2008; Bosze et al., 2020). We found that a splice-blocking MO targeting the intron1-exon2 (i1e2) junction of nascent pcdh18a RNA led with high efficiency to retention of intron 1 in the mRNA (Fig. 2B-a, Fig. S2), resulting in a frameshift past codon 734 and a premature termination codon 13 codons downstream, hence in a protein retaining only 106 out of the 399 amino acids of its cytoplasmic domain (Fig. 2B-b; Fig. S2C). We therefore call this truncated protein Pcdh18a-ΔCP_106_ (Fig. 2B-b). The cytoplasmic domain deleted by the i1e2 MO notably contains the WIRS peptide (Chen et al., 2014) involved in the regulation of F-actin, and the Dab1 interacting domain (Homayouni et al., 2001) that may regulate a tyrosine kinase pathway (Fig. 2B-b), both of which are highly similar to human orthologs (Fig. 2B-c). We then injected this MO in Tg*(pax3a:GFP; TCF:nls-mCherry)* embryos, which labels somite-derived SCPs with GFP and cells of somitic origin with nuclear mCherry (Murayama et al., 2015). By 24-30 hpf, we observed delamination and emigration of SCPs from the ventral side of caudal somites in both control and morphant embryos (Fig. 2C). We then examined how the truncation of the cytoplasmic domain of Pcdh18a occuring in the morphants affected SCP migration. We focused our analysis on the leader cells of the strings of SCPs that emigrated from each caudal somite. We found that the SCP leader cells in pcdh18a-ΔCP_106_ morphants showed a zigzag-like migration pattern (Fig. 2Da-c), and as a result, the straightness of their trajectories was diminished (Fig. 2D-d). Their speed was also affected. We found indeed that the migration of leader SCPs in wild-type embryos can be divided into two phases: a first phase of fast movement lasting about 90 min, followed by a short rest and then a relatively slow second phase (Fig. 2E-a). This velocity change probably corresponds to the transition from migration on the primordial vein surface to a movement in the three-dimensional extracellular matrix further ventrally, synchronized with the migration of and interconnection with endothelial cells in the process of forming the venous plexus (Fig. 1). In contrast, SCP leader cells in morphant embryos migrated at a slower, relatively uniform speed without going through these phases (Fig. 2E-a,b). In addition, in the pcdh18a morphants, migrating SCPs frequently showed a more elongated cell body and thick cellular projections in several directions (Fig. 2Ce,f).

To further analyze the migration behaviour of SCP leader cells in pcdh18a-ΔCP_106_ morphants, we crossed the TgBAC*(cspg4:GAL4)* line with Tg*(UAS:Lifeact-eGFP)* so as to visualize actin dynamics and cellular projections during SCP migration. In the control group, a high-intensity GFP signal was observed in leader cells mainly near the front of the cell – relative to the direction of cell migration (Fig. 3Aa-d, dashed arrows; Movie 3). In contrast, in pcdh18a-ΔCP_106_ morphants, cellular projections were more frequently detected towards the opposite direction (Fig. 3Ae-h; Fig. 3B). A quantitative analysis confirmed that compared to control embryos, leader cells in pcdh18a morphants displayed a similar total number of filopodia (mean=11/cell), as well as towards the direction of migration, but 70% more filopodia than controls towards the opposite direction (Fig. 3C).

**Figure 3.**
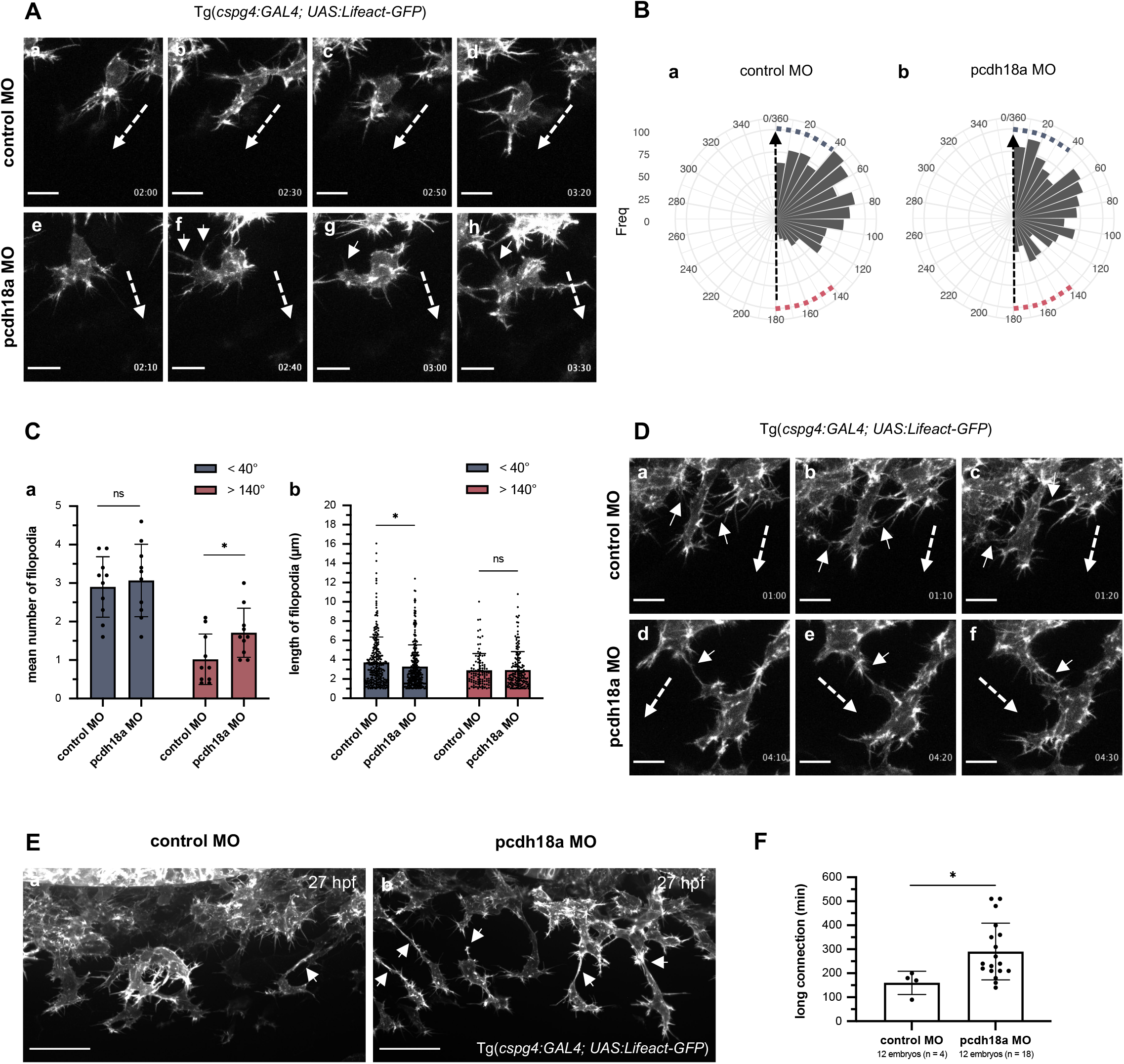
Pcdh18a modulates cell protrusions and connections of migrating SCPs. **(A)** Time-lapse confocal images of individual migrating SCP leader cells from control and pcdh18a-ΔCP_106_ morphant Tg(*cspg4:GAL4; UAS:Lifeact-GFP*) embryos by 26 hpf (see also Movie 3). Images were extracted from movies at 20-30 minutes intervals. Time from the initiation of SCP migration is shown in the lower right of each panel. Dashed and short arrows indicate the direction of SCP migration and filopodia extending in the opposite direction, respectively. Scale bars, 10 µm. **(B)** Quantification of the frequency of filopodia according to their angle relative to the direction of migration in control **(a)** and pcdh18a-ΔCP_106_ **(b)** morphant embryos. In the rose diagram, the dashed arrow indicates the direction of SCP migration; angles < 40° were considered close to the direction of the migration (blue dots); angles > 140° were considered opposite to the direction of migration (red dots). The linear mixed-effect model indicates that pcdh18a-ΔCP_106_ morphants have relative angles increased on average by 14.9° compared to controls (P=0.0041). **(C-a)** Quantification of the mean number of filopodia in sense (<40°, blue) or opposite direction (>140°, red) projected from migrating SCPs in control or pcdh18a-ΔCP_106_ morphant embryos. Each dot represents a biological replicate from 5 independent experiments (n=10 for controls and morphants) (mean±SD; ns, P=0.6666; ^*^, P=0.0284; Student’s t test). **(C-b)** Quantification of the length of filopodia in the sense (blue) or opposite (red) direction of cell migration. Each dot represents a single filopodium (n=392 for controls, 478 for pcdh18a morphants; mean±SD; ns, P=0.5190; ^*^, P=0.0247; Mann-Whitney test). **(D)** Time-lapse confocal acquisitions of migrating leader SCPs from control and morphant Tg(*cspg4:GAL4; UAS:Lifeact-GFP*) embryos around 25-28 hpf. Images were extracted from movies at 10 minutes intervals. In controls, neighboring leader cells interact by brief contact of their filopodia which do not change their direction of migration (a-c, arrows), while in morphants they tend to have longer contact between cells, which can attract and change the trajectory of the neighboring cell (d-f, arrows). Dashed arrows indicate the direction of cell migration. Scale bars,10 µm. **(E)** Confocal spinning disk images of live Tg(*cspg4:GAL4; UAS:Lifeact-GFP*) embryos at 27 hpf. Leader cells tend to show longer connections to follower cells (arrows) in morphants **(b)** compared to controls **(a)**. Scale bars, 30 µm. **(F)** Quantification of the duration of long (>10µm) cell connections between leader and follower cells (Mean±SD; ^*^, P=0.0113; Mann-Whitney test). See also Movie 4.

Next, we focused on the features of cell-cell junctions between leader cells and their followers in the SCP strings emigrating from the somites. At the beginning of their migration in control embryos, the leader cells of neighboring strings sometimes contact each other via filopodia, but these interactions are transient, and they don’t affect the direction of migration of the involved cells (Fig. 3Da-c). In contrast, in pcdh18a-ΔCP_106_ morphants, when such contacts occur, they last longer, with a resulting change of migration direction of at least one of the two cells (Fig. 3Dd-f) - a behaviour that may contribute to the lesser straightness of leader cell trajectories in morphants that we found above (Fig. 2D). In addition, while in WT embryos the leader SCPs tend to separate within the first few hours from the follower SCPs, in pcdh18a-ΔCP_106_ morphants leader cells did not easily separate from follower cells during their migration (Fig. 3E, arrows). Among 12 embryos analyzed by 27 hpf, we found 18 leader cells with a long (>10 µm) connection to the follower cells, and only 4 leader cells with such a long connection in the controls. In addition, these long connections had an almost 2-fold longer mean duration in the morphants (4h50 vs. 2h40, Fig. 3F). Finally, when the connection eventually broke up, it took much more time to retract back to the cell in the morphants (Movie 4).

### Pcdh18a cytoplasmic domain truncation causes a reduction in stromal cell numbers

We next examined whether the lack of most of the cytoplasmic domain of Pcdh18a affected the number of SCPs and their progeny. Cell counting was performed at 26 and 36 hpf, corresponding to the early and latest stages of SCP migration, and at 50 hpf, i.e. once SCs have settled and differentiated within the venous plexus. We found no difference at 26 hpf, but a nearly 2-fold lower stromal cell number in pcdh18a-ΔCP_106_ morphants by 36 hpf and 50 hpf relative to control embryos (Fig. 4A). We therefore quantified the frequency of cell divisions of SCPs/SCs during their migration (24-36 hpf) and settlement/maturation (36-46 hpf) phases, using time-lapse confocal imaging over these two developmental time windows. We found no difference in the number of observed SCP mitoses between control and morphants in the 24-30 hpf and 30-36 hpf intervals (Fig. S3A), and a 2-fold reduction in SCP/SC divisions number in the morphants in the 36-46 hpf interval (Fig. S3B), which matched the nearly 2-fold difference in SCP/SC number at the beginning and end of that time window (Fig. 4A), indicating an actually similar proliferation rate. We therefore quantified the rate of emergence of SCPs from the somites between 26 and 40 hpf, and found a 40% reduction in the morphants (Fig. 4B). Thus, the lower final number of stromal cells in pcdh18a-ΔCP_106_ morphants can be mainly ascribed to a lower number of SCP emergence events from the somites.

**Figure 4.**
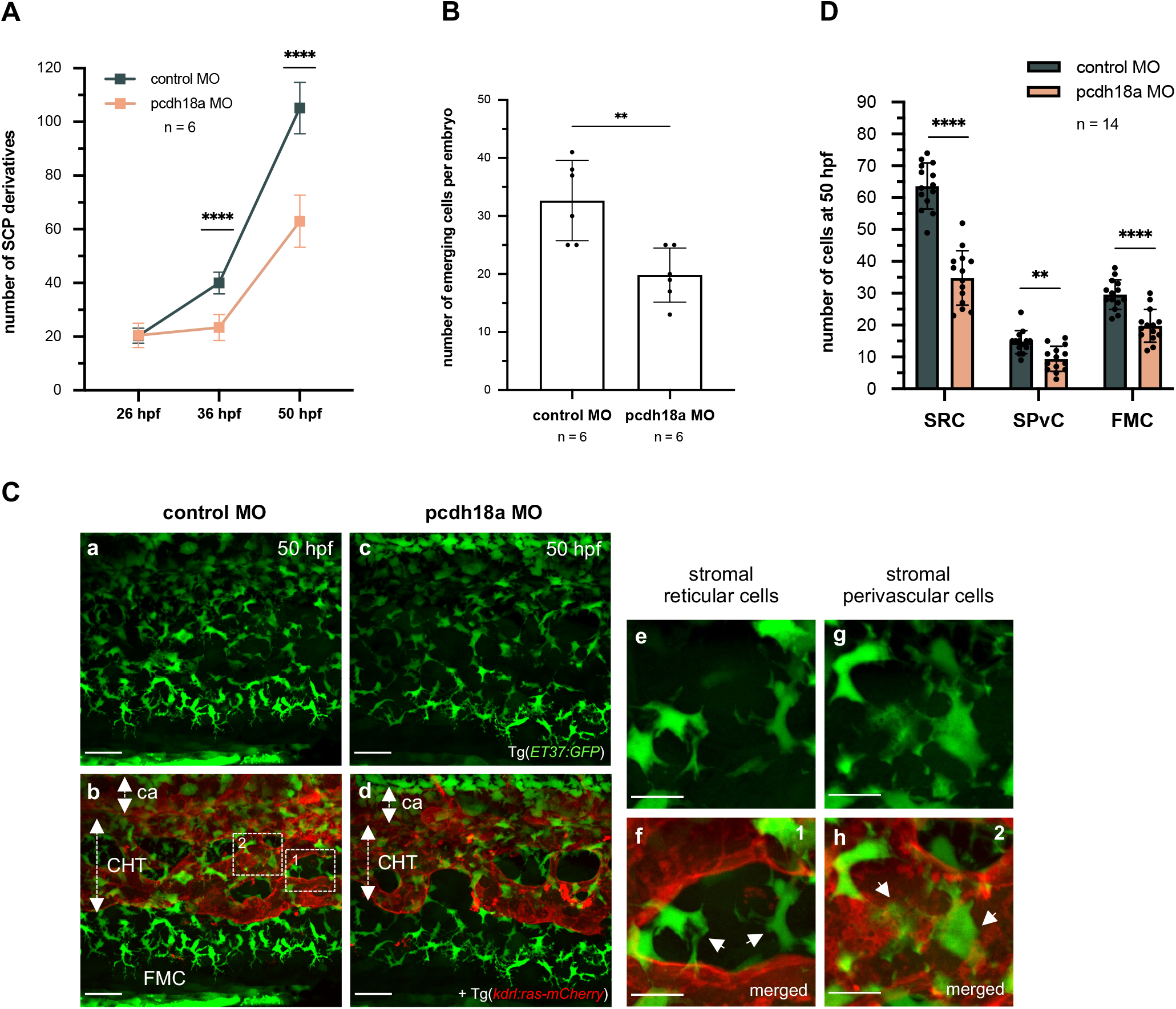
Pcdh18a C-terminal truncation reduces the number of stromal cells. **(A)** Counting of ET37:GFP+ SCP derivatives in the developing ventral tail over a 5 somites width at 26, 36 and 50 hpf. Plots represent biological replicates from a single experiment; n=6 for control and pcdh18a-ΔCP_106_ morphants (Mean±SD; ^****^, P<0.0001, Student’s t test). **(B)** Counts of the number of emerging SCPs within a 5 somites width, from 24 to 40 hpf (from the same time-lapse confocal sequences as in Fig. 2C), n=6 embryos per condition (mean±SD; ^**^, P=0.0037; Student’s t test). **(C) (a-d)** Confocal spinning disk projections of live Tg(*ET37:GFP; kdrl:ras-mCherry*) control or pcdh18a-ΔCP_106_ embryos at 50 hpf. ca, caudal artery; FMC, fin mesenchymal cells. Scale bars, 30 µm. Dotted frames in **b** indicate regions enlarged in **(e-h)**, which show the two stromal cell types found in the CHT at 50 hpf. Stromal reticular cells (SRCs) interconnect the vascular segments of the plexus and seem to maintain its architecture (**e** and arrows in **f**), whereas stromal perivascular cells (SPvC) are spread along and around the vessels (**g**, and arrows in **h**). Scale bars, 10 µm. **(D)** Numbers of the three types of SCP derivatives – SRCs and SPvCs in the CHT, and FMCs in the caudal fin in 14 embryos per condition, counted over a 5 somites width (mean±SD; ^**^, P=0.0011; ^****^, P<0.0001; Student’s t test).

By 50 hpf, we could identify two subtypes of SCP-derived stromal cells settled in the CHT niche (Fig. 4C) - stromal reticular cells (SRCs), which extend cellular projections that interconnect the vascular segments of the venous plexus (Fig. 4C-a,b,e,f), and stromal perivascular cells (SPvCs), that adhere to endothelial cells of the plexus over their entire surface (Fig. 4C-a,b,g,h). We thus compared the numbers of SCP-derived SRCs, SPvCs and FMCs between pcdh18a-ΔCP_106_ morphants and controls. All three cell types showed a similar substantial decrease in the morphants (45% for SRCs, 36% for SPvCs, and 33% for FMCs; Fig. 4C,D).

### Pcdh18a cytoplasmic domain conditions stromal cells guidance of venous plexus morphogenesis

We next investigated how truncation of the Pcdh18a cytoplasmic domain affected the behaviour of SCPs relative to the process of venous plexus formation, using time-lapse imaging of Tg*(pax3a:GFP; kdrl:ras-mCherry)* embryos between 26 and 36 hpf (Fig. 5A; Movie 5). In the control group, SCPs were always present at the forefront of endothelial cells that actively branched and migrated to the ventral side, and they physically interacted with them via cellular projections/filopodia (Fig. 5Aa-d). In contrast, while both endothelial cells and SCPs migrated to the ventral side in pcdh18a-ΔCP_106_ morphants, they often did not interact (Fig. 5Ae-h; Movie 5). In addition, while in control embryos SCPs were evenly distributed among migrating endothelial cells, and at some distance from each other (Fig. 5Aa-d), leading SCPs in morphant embryos were often clustered at the forefront between migrating endothelial cells (Fig. 5Ae-h).

**Figure 5.**
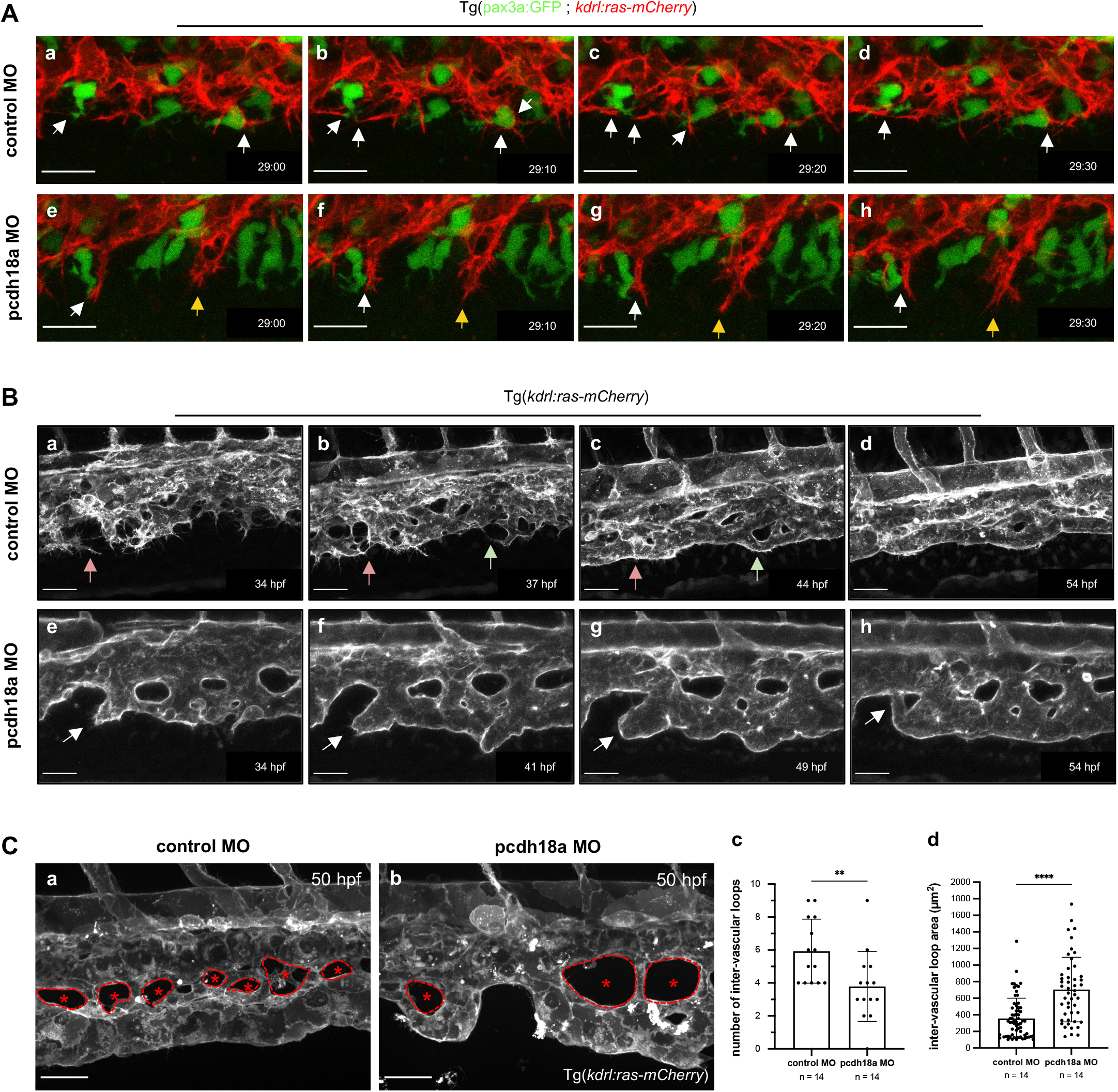
Pcdh18a truncation impacts on venous plexus morphogenesis. **(A)** Time-lapse confocal acquisitions from control **(a-d)** and pcdh18a-ΔCP_106_morphant **(e-h)** Tg(*pax3a:GFP; kdrl:ras-mCherry*) embryos (see also Movie 5). Confocal projections at the ventral part of the primordial vein during active sprouting angiogenesis and associated co-migration of stromal cells. Images were extracted from movies at intervals of 10 minutes from 29 hpf to 29.5 hpf. Dynamic interactions between stromal and endothelial cells (white arrows) were frequently observed in control embryos whereas pcdh18a morphant displayed less interactions and vascular endothelial cells actively migrated to the ventral side regardless of the involvement of stromal cells (yellow arrows). Scale bars, 20 µm. **(B)** Time-lapse confocal images from control **(a-d)** and pcdh18a **(e-h)** MO-injected Tg(*kdrl:ras-mCherry*) embryos at later stages (34-54 hpf). See also Movie 6. Arrows in two different colors follow the formation of new vascular segments. Unusual tubulogenesis was frequently observed in pcdh18a morphants at the most ventral side of the plexus where the definitive caudal vein forms (white arrows). Scale bars, 30 µm. **(C)** Morphological differences in the formation of intervascular loops (IVLs). **(a**,**b)** Confocal spinning disk acquisitions of live transgenic Tg(*kdrl:ras-mCherry*) control or pcdh18a MO-injected embryos at 50 hpf. IVLs are indicated by asterisks and the border of each IVL is delineated by a dashed line. Scale bars, 30 µm. **(c**,**d)** Number **(c)** and surface in µm^2^ **(d)** of IVLs over a 5 somites width in controls and morphants; n=14 embryos per condition, from 2 independent experiments (mean±SD; ^**^, P=0.0066; ^****^, P<0.0001; Mann-Whitney test).

We then focused on the subsequent process of venous plexus remodeling (34-54 hpf). Whereas in control embryos, endothelial cells forming the plexus still extended numerous filopodia and projections (Fig. 5B,a-d; Movie 6), in the morphants they no longer did so, and rather appeared to have already fused more extensively with each other (Fig. 5B,e-h; Movie 6). The resulting plexus had fewer branching points compared with controls and tended to show a simpler/flatter structure, with fewer but larger inter-vascular loops (IVLs), and a frequent malformation of the resulting definitive caudal vein, caused by improper fusion events deviating it from its normal course parallel to the body axis (Fig. 5Be-h, arrows; Fig. 5Ca,b). A quantitative analysis confirmed that the number of IVLs was decreased by one third relative to controls, whereas their mean 2D surface area was increased 2-fold (Fig. 5Cc,d).

### Pcdh18a cytoplasmic truncation leads to excess deposition of fibronectin in the future hematopoietic niche

Since truncation of the cytoplasmic domain of Pcdh18a affected not only SCP numbers and migration behaviour, but also their interaction with endothelial cells and the final structure of the venous plexus, we speculated that in addition to a possibly direct involvement of Pcdh18a in these processes, there may be other factors affected by Pcdh18a truncation that may act on both SCPs and vascular endothelial cells in terms of their interactions during venous plexus morphogenesis. Therefore, we looked for genes encoding extracellular matrix molecules expressed in the venous plexus area during its formation (24-36 hpf). qPCR analysis of *syndecan2, cspg4* (chondroitin sulfate proteoglycan 4), *hspg2* (heparan sulfate proteoglycan 2), *fn1a* and *fn1b* (fibronectin 1a and 1b) expression in the tails of control and pcdh18a-ΔCP_106_ morphants at 36 hpf revealed that *fn1a* and more so *fn1b* were upregulated in the morphants (Fig. S4A). Immunostaining revealed that fibronectin was highly deposited at somite boundaries and the venous plexus in pcdh18a-ΔCP_106_morphants (Fig. S4B). Optical transverse sections of pcdh18a-ΔCP_106_ morphants (Fig. S4Bg-i) showed the surface labeling of mCherry+ endothelial cells by the anti-Fibronectin antibody. We then performed WISH analysis to identify the cells that express *fn1b*. We found it expressed in caudal somites by 23 hpf, and strongly up-regulated there in pcdh18a-ΔCP_106_ morphants (Fig. S4Ca,b). At 26 hpf, specific expression of *fn1b* was observed in the endothelial sprouts that will form the venous plexus of morphant but not control embryos (Fig. S4C,c-d’).Then *fn1b* expression was no longer detected at 36 hpf in the tail of control embryos, but still strongly present in the morphants (Fig. S4C-e,f).

### Pcdh18a cytoplasmic truncation leads to a non-functional hematopoietic niche

We finally examined to what extent Pcdh18a cytoplasmic truncation affected the resulting niche function of the CHT for HSPCs. We found that in pcdh18a-ΔCP_106_ morphants, definitive aorta-derived HSPCs expressing *myb* were present in the sub-aortic space of the trunk by 36 hpf in numbers similar to control embryos (Fig. 6A-a,b). Yet by 2.5 and 3 dpf, only few of them had homed to the CHT in the morphants (as evidenced by CD41-GFP^low^ HSPCs and CD41-GFP^high^ pro-thrombocytes, Fig. 6B, and by *myb* expression, Fig. 6A-c,d). The incapacity of the morphant CHT to support hematopoiesis became even more evident by 5 dpf, when HSPC numbers in this niche reach their maximum in wild-type larvae (Murayama et al., 2006) (Fig. 6A-e,f).

**Figure 6.**
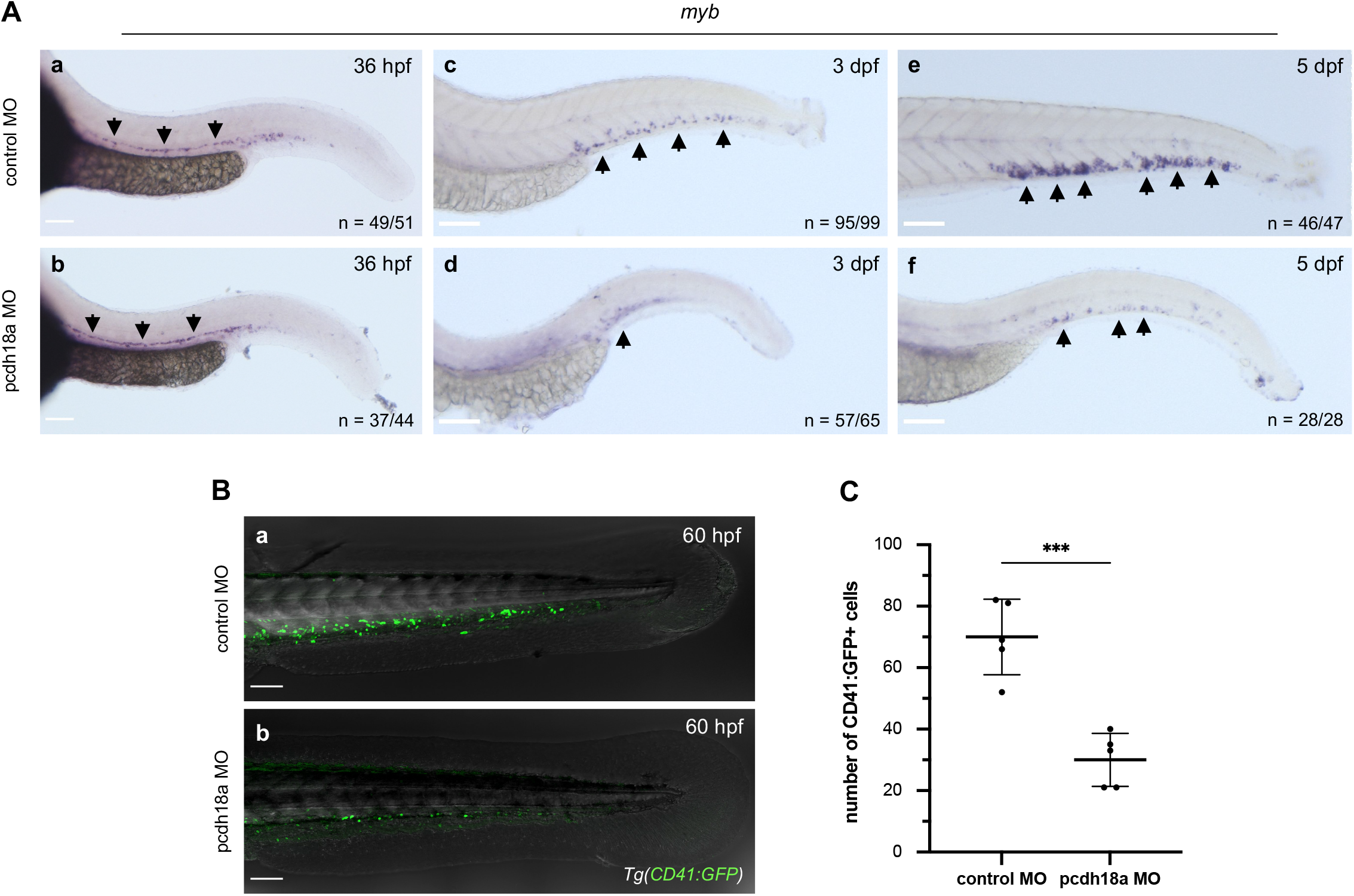
The Pcdh18a cytoplasmic domain is required for HSPC lodgment and expansion in the CHT. **(A)** Whole-mount in situ hybridization for *myb* expression on control or pcdh18a-ΔCP_106_ morphant embryos. Scale bars, 100 µm. **(a-b)** At 36 hpf, control and pcdh18a-ΔCP_106_ morphants showed similar *myb* signals along the dorsal aorta (arrows), meaning they can produce definitive HSPCs properly. **(c-f)** Pcdh18a-ΔCP_106_ morphants showed a lower number of *myb+* cells signal in their CHT than controls at 3 dpf, and even more so at 5 dpf. **(B-a**,**b)** Confocal acquisitions of live Tg(*CD41:GFP*) control or pcdh18a-ΔCP_106_ morphant embryos at 60 hpf. Scale bars, 100 µm. **(C)** Quantification of the number of CD41:GFP+ cells in the CHT of controls or morphants at 60 hpf (n=5 embryos per condition; mean±SD; ^***^, P=0.0003; Student’s t-test).

### Adhesion to endothelium via the extracellular domain of Pcdh18a determines stromal cell subtypes

In addition to cell-cell adhesion through homophilic binding via their extracellular domain (ECD) (Morishita and Yagi, 2007), protocadherins may also engage in heterophilic interactions (Jontes, 2016). We therefore wondered if the ECD of Pcdh18a might be directly involved in the adhesion of SCPs to vascular endothelial cells. We first searched for cell adhesion motifs such as RGD in the ECD of Pcdh18a. Instead of the RGD motif, we found there two REDV motifs, that were originally identified in human plasma fibronectin as a peptide that binds to vascular endothelial cells (Fig. 7A-a). Therefore, we decided to investigate how SCPs behave when a mutation is introduced into the REDV motifs in Pcdh18a. Firstly, to choose the substitution patterns of the four amino acid residues of REDV, we used the SWISS-MODEL server for protein structure prediction to see how various mutations may affect the conformation of the ECD of Pcdh18a. We performed structural predictions for four different substitution patterns - AAAA, AEAV, AEDA and RAAV - and found that the substitution from REDV to AEAV best retained the three-dimensional structure of Pcdh18a, at least under the condition of homophilic interaction (Fig. 7A-a,b).

**Figure 7.**
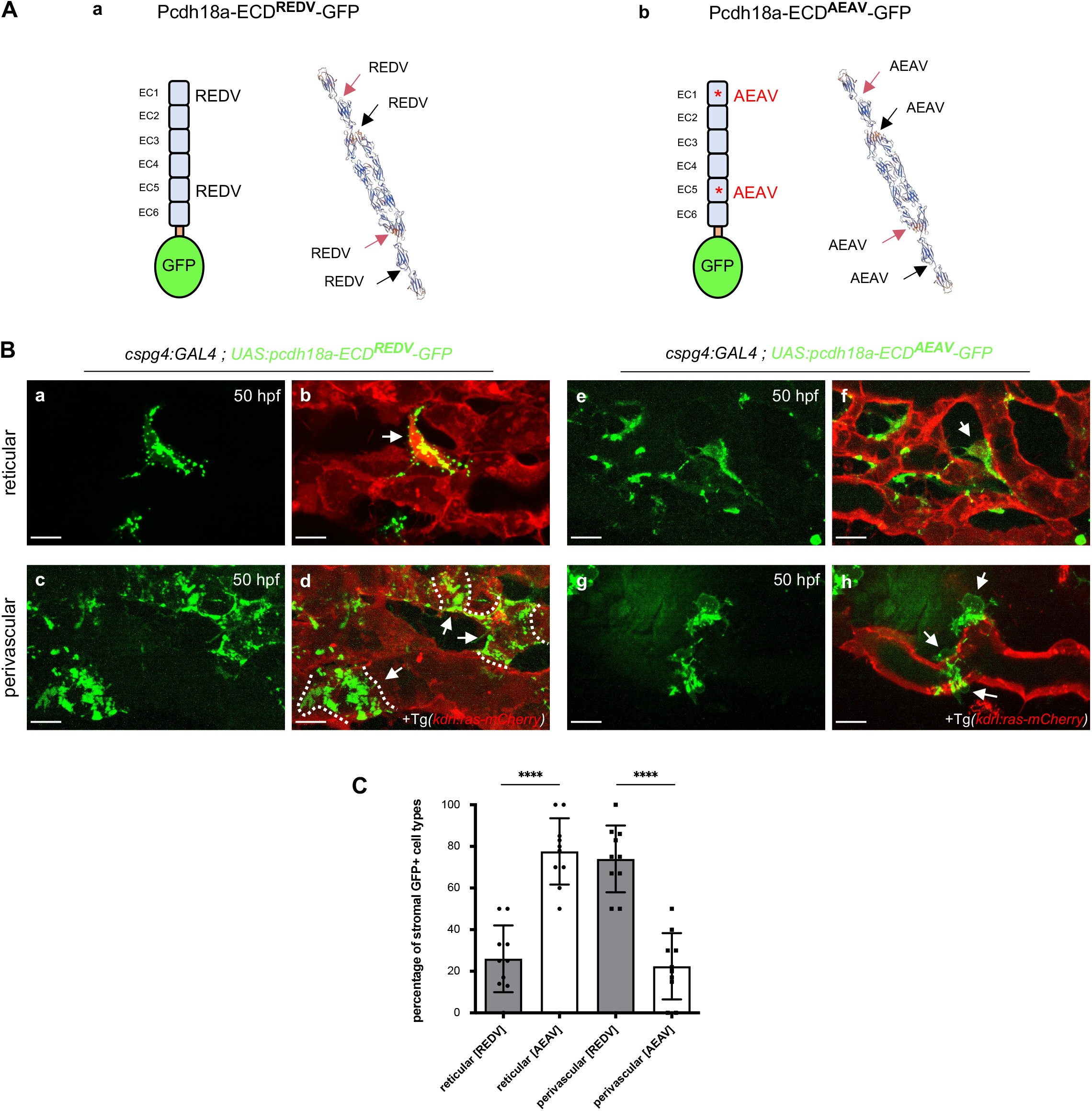
The extracellular domain of Pcdh18a mediates the adhesion of stromal cells to endothelial cells. **(A)** Fusion proteins expressed in SCPs via the Gal4/UAS system, with the ICD of Pcdh18a substituted by GFP, and tertiary structure prediction for WT and mutant forms of the extracellular domain (ECD), using SWISS-MODEL (expasy.org). Red and black arrows point at the two REDV (WT) or AEAV (mutant) motifs in each of two ECDs in homophilic trans interaction. **(B)** Confocal spinning disk projections of live Tg(*cspg4:GAL4; UAS:pcdh18a-ECD*^*REDV*^ or ^*AEAV*^*-GFP; kdrl:ras-mCherry*) embryos in the caudal venous plexus at 50 hpf. **(a-d)** Pcdh18a-ECD^REDV^-GFP expressing reticular (a,b) or perivascular (c,d) stromal cells (green) overlaid on the endothelial cells (red). Dashed lines indicate the border of each SPvC. **(e-h)** Pcdh18a-ECD^AEAV^-GFP expressing reticular (e,f) or perivascular (g,h) stromal cells (green) overlaid on the endothelial cells (red). GFP+ SPvC typically showed an only partial adherence to the endothelium. Scale bars, 10 µm. **(C)** Histogram showing the percentage of reticular vs. perivascular GFP+ stromal cells observed among the pcdh18a-ECD^AEAV^-GFP or pcdh18a-ECD^REDV^-GFP expressing stromal cells at 50 hpf. 10 embryos of each condition were analyzed (cell counting was performed by dividing the CHT into three parts to cover a 5 somites width) from a single experiment (mean±SD; ^****^, P< 0.0001; Student’s t-test).

Next, we analyzed the effect of these amino-acid changes *in vivo* by expressing Pcdh18a carrying wild-type (REDV) or mutant (AEAV) sequences, fused to GFP instead of Pcdh18a intracellular domain. To overexpress either form specifically in SCPs, we inserted these constructs downstream of a UAS promoter (UAS:pcdh18a-ECD^REDV^-GFP and UAS:pcdh18a-ECD^AEAV^-GFP), and injected the resulting plasmids into Tg*(cspg4:Gal4; kdrl:ras-mCherry)* embryos, so as to also visualize vascular endothelial cells. We then analyzed the resulting embryos by live confocal imaging at 50 hpf, i.e. when the SCPs have differentiated into stromal cell subtypes. We found that in embryos overexpressing the WT (REDV) Pcdh18a ECD, GFP+ stromal cells were mainly of the perivascular subtype, i.e. closely adherent to vascular endothelial cells of the venous plexus (Fig. 7Bc,d). In contrast, GFP+ stromal cells overexpressing the ECD of Pcdh18a into which the AEAV mutation was introduced were mainly of the reticular type (Fig. 7Be,f), and those that were more perivascular displayed a characteristic defect in attachment to the vascular endothelium. They were generally smaller in 2D size than those of control embryos, and only part of the cell adhered to the endothelium (Fig. 7Bg,h).

We then counted the proportions of stromal cells of the two subtypes - reticular and perivascular - among GFP+ cells overexpressing either the WT or the mutant Pcdh18a ECD form (Fig. 7C). While in normal uninjected embryos, the ratio of stromal reticular to perivascular cell number is 80:20 (Fig. 4D), overexpression in stromal cells of the WT Pcdh18a ECD form led to a sheer inversion of this ratio, to 20:80, indicating that this overexpression strongly favored the perivascular fate. In striking contrast, overexpression in stromal cells of the mutated (AEAV) Pcdh18a ECD form led to a reticular versus perivascular ratio similar to that found in uninjected embryos, i.e. 80:20. Altogether these qualitative and quantitative results demonstrate that the ECD of Pcdh18a is required for the extensive adhesion of stromal cells to endothelial cells that characterizes the perivascular subtype, and that this effect is strictly dependent on the REDV peptides of the Pcdh18a ECD.

## Discussion

Stromal cell progenitors (SCPs) differentiate from zebrafish caudal somites via an epithelial to mesenchymal transition (EMT). The timing of EMT in SCPs during somite development is still uncertain, but probably SCPs remain in the cluster for a while after the EMT, then delaminate ventrally around 23-24 hpf. Mesenchymal cell migration is characterized by cell polarization, forming a leading edge that extends actin-rich protrusions, and subsequent migration by retraction of the contractile rear. Cell migration and adhesions are interdependent processes that influence diverse cellular fates. Neural crest cells (NCCs), a highly motile, EMT-derived cell population, undergo collective cell migration, and the involvement of cadherins in this process is well-known. Like NCCs, SCPs undergo collective migration, but we call this a ‘semi-collective migration’ because the number of cells in a group is fewer (2-3 cells) than for NCCs. Semi-collective migration of SCPs occurs immediately after delamination from the SCP clusters at the ventral side of somites, and shortly thereafter, the leader cell releases itself from the follower. This characteristic suggests that binding through cell adhesion molecules among SCPs may be less tight (or more flexible) than that of NCCs. We observed no expression of cadherin genes in the SCPs, suggesting that protocadherins such as Pcdh18a may act as a substitute for cadherins in SCPs. Pcdh18a is indeed expressed in the SCPs within the clusters and during their subsequent migration. We found that it facilitates their overall emergence from the clusters over time, since a truncation of its ICD caused a nearly 2-fold reduction in the total number of SCPs that emerged from the somites, and it also promotes their ventralwards navigation, as the latter is less direct when Pcdh18a ICD is truncated. This may be connected to the fact that the truncated Pcdh18a no longer contains the WIRS sequence that can bind the WAVE complex and thereby trigger actin polymerization. We also found that the Pcdh18a ICD is involved in repulsion between leader cells among migrating SCPs. This is reminiscent of previous data about two other δ 2-Pcdhs. A study based on cell aggregation assays found that Pcdh19 lacking its ICD induced the formation of larger aggregates than full-length Pcdh19 (Tai et al., 2010). Then two studies reported repulsive forces among axons (Hayashi et al., 2014), and among abducens motor neurons in zebrafish (Asakawa and Kawakami, 2018), that were dependent on the ICD of Pcdh17. Thus, it appears that the ICD of δ2-Pcdhs can have an adhesion-inhibiting role, although the mechanism is still unknown.

The presence of Pcdhs, which mediate weaker homophilic interactions than cadherins, in migrating SCPs, may help ensure the flexibility to switch from the collective migration of SCPs to their interaction with vascular endothelial cells in a short time after delaminating from somite clusters. The endothelial cells forming the caudal venous plexus express no Pcdh18, but a variety of integrins (Xue et al., 2019), notably α5β1 that can bind to the RGD motif present on other cells or proteins of the ICM. While the ECD of Pcdh18a contains no RGD motif, we found in it two copies of a REDV motif that had been reported to mediate protein binding to endothelium expressing integrin α4β1, and these two motifs were in exposed position on the predicted 3D structure of Pcdh18a. By expressing a WT or mutated form of Pcdh18a specifically in SCPs/SCs, we demonstrated that the ECD of Pcdh18a is essential, via its REDV motifs, for the adhesion of SCPs/SCs to the CVP endothelium. This is probably the first documented case of trans heterophilic binding of a δ 2-family protocadherin in vivo, and it bears a high biological significance, as it conditions the formation of a functional hematopoietic niche (CHT). Furthermore, overexpression of Pcdh18a-ECD at the surface of SCPs/SCs strongly favored the installment of SCPs as perivascular rather than reticular stromal cells, which normally largely predominate. It remains to be explored whether this shift is accompanied by actual changes in differentiation at the molecular level.

In addition to its ECD, the ICD of Pcdh18a also appears to be important for the proper interactions of SCPs/SCs and endothelial cells. Indeed pcdh18a-ΔCP_106_morphants display morphological defects of the final venous plexus. SCPs may orchestrate the migration and orderly fusion pattern of endothelial cells to form a venous plexus of adequate shape, possibly by refraining them to fuse too rapidly and extensively - as seems to happen in the pcdh18a-ΔCP_106_ morphants. We note that SCPs are distributed quite evenly among endothelial cells in the process of forming the plexus, thus maximizing the number of endothelial cells in contact with at least one SCP, whereas SCPs are both nearly 2-fold less numerous and less evenly distributed in pcdh18a-ΔCP_106_morphants, which may compromise this orchestrating function.

We also considered the possibility that changes in the ECM may be involved in the abnormal remodeling of the venous plexus in pcdh18a-ΔCP_106_ morphants. Surprisingly, we found that fibronectin was excessively expressed and accumulated in the CHT of these morphants. WISH analysis showed that *fn1b* expression was markedly elevated in the caudal somites of pcdh18a-ΔCP_106_ morphants by 23 hpf, and then expressed in the caudal vasculature by 26 hpf, unlike in control embryos. This excess accumulation of fibronectin in the caudal region of pcdh18a-ΔCP_106_ morphants may contribute to the observed defects in venous plexus formation. Zebrafish fibronectin 1b contains no REDV motif, but two RGD motifs and two LDV motifs, that can bind to integrins α5β1 and α4β1 (Komoriya et al., 1991), respectively, and both of these integrins are expressed by endothelial cells forming the venous plexus (Xue et al., 2019). Thus, excess fibronectin 1b may compete with the binding of SCPs/SCs to endothelial cells of the venous plexus, notably through its LDV motifs, which bind the same α4β1 integrin as the REDV motifs of Pcdh18a-ECD that we found to mediate stromal cell binding to these endothelial cells. It remains unclear why and how a deficiency in the Pcdh18a-ICD causes an increase in expression of the fibronectin genes in caudal somites and endothelial cells. We speculate that unusual cell-cell interactions caused by Pcdh18a truncation in SCPs may induce a microenvironmental stress and promote the secretion of extracellular matrix, notably fibronectin, by nearby cells.

This study adds new insights into the functions of Pcdh18a, as an adhesion molecule that plays an essential role in the development of SCPs, a newly recognized cell model arising from an EMT. After the EMT, 1) SCPs remain in the clusters at the ventral side of somites for a while, 2) then undergo semi-collective cell migration in which 2-3 cells from a string, but shortly after, 3) switch binding partners from SCPs to vascular endothelial cells. Pcdh18a is continuously expressed in SCPs during these three steps. Thus, in a different state of cell dynamics in SCPs, Pcdh18a may flexibly regulate the functions of the extracellular and intracellular domains in response to the microenvironment. Substrate selectivity or affinity in mesenchymal cell migration is also a crucial point in the field of cancer research. Cancer cells are known to disseminate in amoeboid, mesenchymal, and collective cell migration modes, which are affected by tumor microenvironment and treatment (Wolf et al., 2003). Recent discussions have raised the possibility that such a plasticity of cancer cell migration is a hindrance to certain treatments (reviewed in Wu et al., 2021; Clark and Vignjevic, 2015). Our research contributes to the understanding of cell behavior *in vivo* by analyzing the function of a protein that could be a molecular switch of ‘migration modes’ affecting cell plasticity during cancer metastasis and tumorigenesis.

## Materials and Methods

### Zebrafish husbandry

Fish maintenance at the Institute follows the regulation of the 2010/63 UE European directives. We used several zebrafish transgenic lines, including Tg(*pax3a:eGFP*) (Seger et al., 2011), Tg(*ET37:eGFP*) (Parinov et al., 2004), Tg(*kdrl:ras-mCherry*) (Chi et al., 2008), Tg(*UAS;Lifeact-eGFP*) (Helker et al., 2013), Tg(*cspg4:Gal; UAS:RFP*) created in the lab. Embryos were collected and raised at 28°C, or at 24°C, taking in account the 25% delay in development relative to 28°C, as described (Kimmel et al., 1995). In order to prevent pigmentation, embryos develop in N-Phenylthiourea (PTU, *Sigma-Aldrich*, P7629) in Volvic source water (0.003% final), supplemented with 280 mg/L methylene blue (*Sigma-Aldrich*, M4159). For experiments, embryos were anesthetized in tricaine 1x (*Sigma-Aldrich*, A5040), equivalent to a final concentration of 160 µg/ml.

### Plasmid constructs

First, we cloned the *pcdh18a* whole sequence in a pG1_Tol2_hsp70 vector, from cDNA made from a pool of 25 to 36 hpf embryo tails (see section mRNA extraction and cDNA synthesis). We separated the gene in 4 fragments (around 1000 bp each) and amplified them by using the Expand High Fidelity Polymerase (*Sigma-Aldrich*, 11732641001) and the primers 1 to 8 (see Table 1). We used the TA cloning kit (*Thermo-Fisher*, K202020) to clone the DNA fragments, then the Gibson Assembly strategy (*NEB*, E2611S) to obtain the whole gene sequence, by using primers 9 to 16 (see Table 1). To generate the UAS:pcdh18aECD^REDV/AEAV^-GFP constructs, we used the pG1_Tol2_hsp70:pcdh18a as a template and the Gibson Assembly strategy with the vector Tol2_4xnr_UAS_eGFP and the primers 17 to 20 for the REDV (non-mutated) form and the intermediate primers 21 to 24 for the AEAV (mutated) form by using a PCR-directed-mutagenesis approach (see Table 1, mutated sites in bold). The constructs were fully verified by sequencing.

**Table 1.**
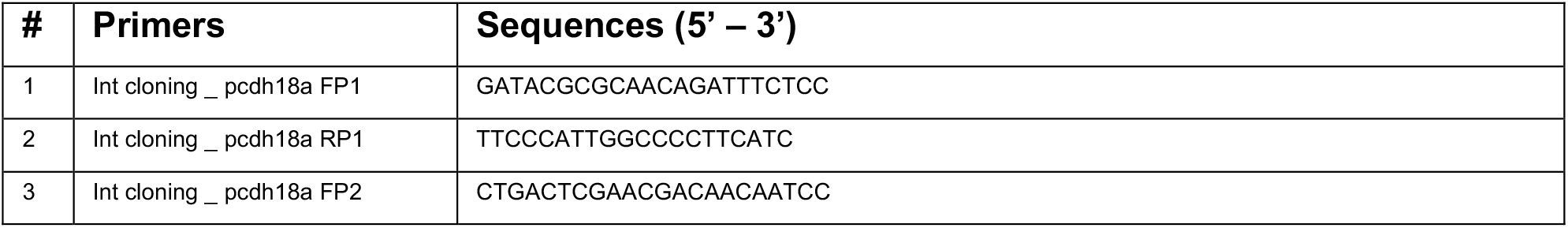
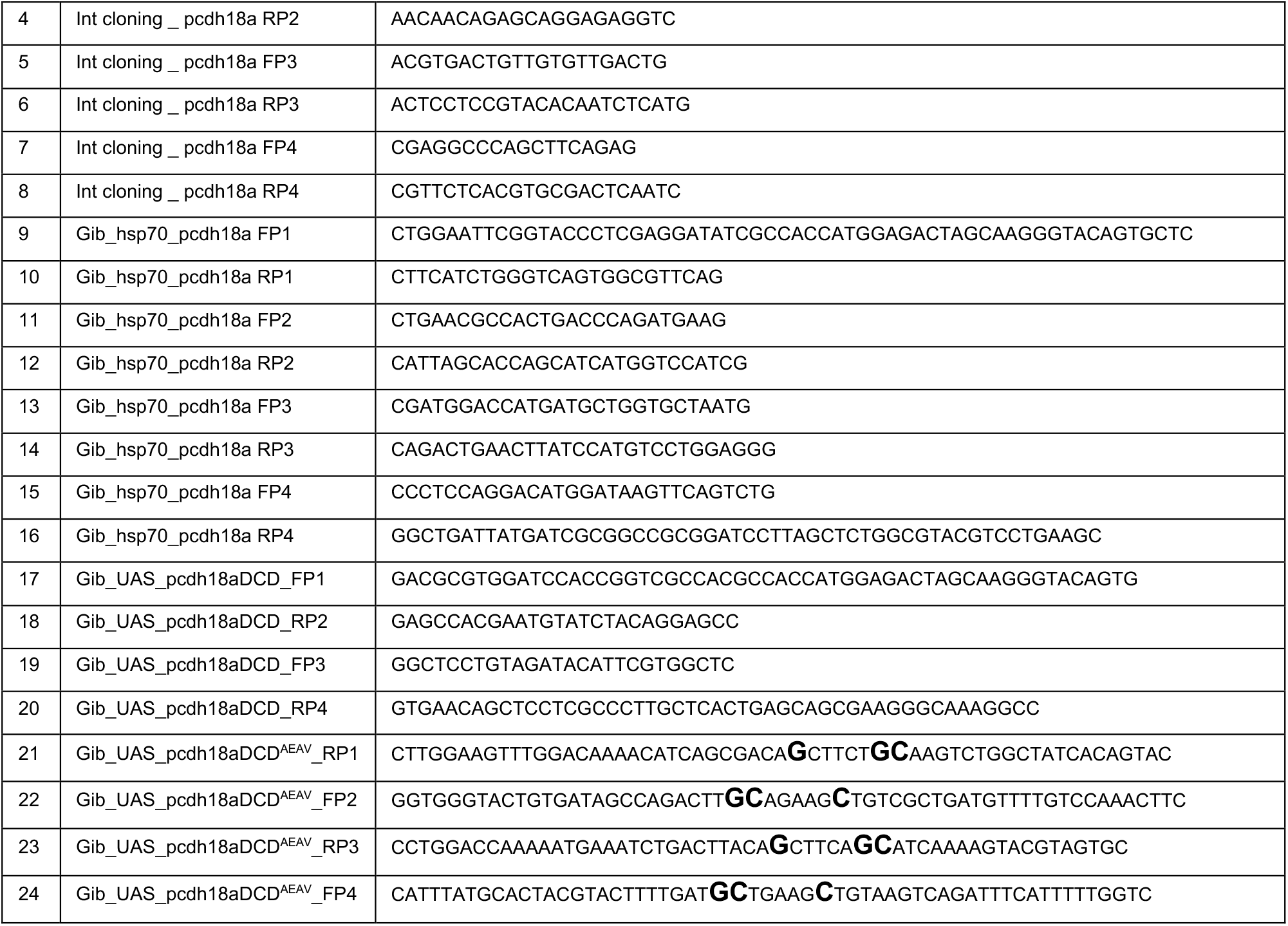
Primers used for intermediate and Gibson cloning of *pcdh18a*.

### Plasmid and morpholino injections

Plasmid constructs were purified using the NucleoBond Xtra Midi endotoxin free kit (*Macherey Nagel*, 740420.10). We co-injected into one-cell stage zebrafish embryos 15 pg of UAS:pcdh18aDCD^REDV/AEAV^-GFP construct with 25 pg of tol2 transposase mRNA, transcribed from linearized pCS-zT2TP plasmid (Suster et al., 2011) using the mMESSAGE mMACHINE SP6 kit (*Ambion*, AM1340). Embryos expressing GFP were sorted and imaged at the desired stages.

A splice blocking morpholino targeting the intron1-exon2 splice junction of *pcdh18a* (5’ – TACTGACCCTGATGGAGTATTGAGA – 3’), was injected at an amount of 4 ng into one-cell-stage zebrafish embryos. A standard control Mo from GeneTools (5’ – CCTCTTACCTCAGTTACAATTTATA – 3’) targeting a mutated form of the human β-globin pre-mRNA was injected at the same amount of 4 ng in control embryos. Control and pcdh18 i1-e2 morphant embryos were picked randomly for fluorescence microscopy.

### Whole-mount immunohistochemistry

Dechorionated and anesthetized embryos were fixed at desired stages in 4% methanol-free formaldehyde (FA) (*Polysciences*, 040181) during 2 hours at RT and then rinsed in PBSDT (PBS 1x, 0.01% DMSO, 0.1% Triton x100 (*Sigma*, T9284)) for use in the week. The first day, embryos were treated with Tween 0.1% during 30 min at RT, then with acetone during 20 min at -20°C, rinsed in HBSS 1x (*Invitrogen*, Gibco 14025), supplemented with 0.1% Tween, then treated with collagenase (*Sigma*, C9891) at a final concentration of 1 μg/ml, rinsed in PBSDT and incubated in blocking solution 1x WBR (*Roche*, 11921673001) for at least 4 hours. They were then incubated overnight at 4°C with primary antibodies – chicken anti-GFP (*Abcam*, ab13970, used at 1/800), mouse anti-mCherry (*Abcam*, ab125096, used at 1/100), rabbit anti-fibronectin (*Sigma-Aldrich*, F3648, used at 1/100). On the second day, embryos were rinsed several times in PBSDT and endogenous peroxydase was inactivated by a treatment with H2O2 6% (*Sigma*, H1009) during 30 min. Then embryos were incubated in the blocking solution for at least 4 hours and incubated overnight at 4°C with secondary antibodies anti-chicken-AF488 (*Invitrogen*, A11039, used at 1/800), anti-rabbit-HRP (*ThermoFisher Scientific*, G-21234, used at 1/300), anti-mouse-HRP (*ThermoFisher Scientific*, G-21040, used at 1/300). The last day, samples were rinsed several times in PBT and treated with imidazole 1M (*Sigma*, I5513)/PBT supplemented with H2O2, Alexa Fluor™ 546 Tyramide Reagent (*Invitrogen*, B40954, used at 1/200) or Alexa Fluor™ 647 Tyramide Reagent (*Invitrogen*, B40958, used at 1/25). Finally, embryos were treated with DAPI (*Thermo Scientific*, 62248) and gradually transfered to glycerol 80%.

### Whole-mount in situ hybridization

Dechorionated and anesthetized embryos were fixed at desired stages in 4% methanol-free FA (*Polysciences*, 040181) overnight at 4°C and stored in 100% methanol at -20°C. The protocol used is based on (Thisse and Thisse, 2008). The first day, embryos are progressively rehydrated in PBT and permeabilized with 10 mg/ml Proteinase K (*Ambion*, AM2546) for 5 min, then refixed in 4% FA for 20 min. Embryos are incubated for at least 4 hours at 60°C in pre-hybridization buffer (HB+), then with 200 ng of Digoxigenin-labeled probe. Probes are synthetized from a PCR product or a linearized plasmid (see Table 2) with the T7 or SP6 RNA polymerase (*Promega*, P4074 or P4084) in presence of DIG-UTP nucleotides. The second day, embryos are rinsed in HB-buffer (HB+ without torula RNA and heparin) supplemented with saline-sodium citrate buffer (SSC, *Sigma*, S6639) and then transferred gradually to PBT. They are incubated for at least 4 hours in blocking solution 1x WBR (*Roche*, 11921673001) and incubated overnight at 4°C with anti-Digoxigenin antibody coupled to alkaline phosphatase (*Roche*, 11207733910, used at 1/4000). The last day, embryos are rinsed several times in PBT and the staining reaction is performed. Samples are incubated in NTMT buffer (100 mM Tris HCl pH 9.5, 50 mM MgCl2, 100 mM NaCl, 0.1% Tween) for at least 5 min and then with NBT/BCIP (*Sigma*, N6639, B-8503) diluted in the NTMT buffer. The reaction is stopped by replacing the solution with PBT, and embryos are stored in 80% glycerol. Bright-field images are captured using a ZEISS Axio Zoom.V16 with a 160x zoom.

**Table 2.**
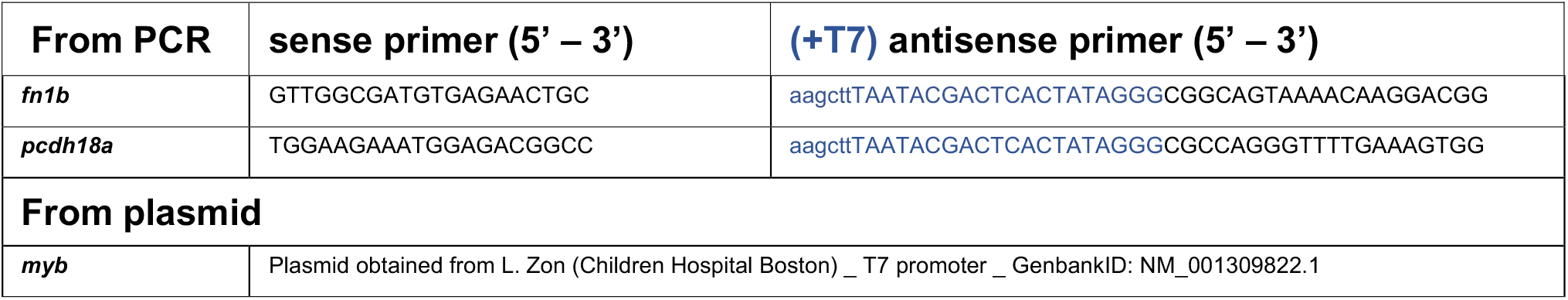
Primers and plasmids used to synthesize WISH probes.

### RNA extraction and cDNA synthesis

We anesthetized embryos and cut their tails just past the yolk extension at desired stages. Batches of 10 tails were immediately frozen in liquid nitrogen and stored at - 80°C. 30 to 60 tails were pooled per biological replicate and 3 biological replicates were used per condition.

For mRNA extraction we used TRIzol™ (*Invitrogen*, 15596026) and lysed tissues by mechanical pipetting. A volume of 1:5 chloroform was added, samples were vortexed and centrifuged. The aqueous phase was transferred to a new tube and a volume of 1:1 isopropanol was added, samples were incubated 30 min at -20°C and centrifuged. The pellet was rinsed in 75% ethanol and resuspended in RNase free water. Samples were treated with DNase I (*Invitrogen*, AM2239), precipitated a second time with GlycoBlue™ (*Invitrogen*, AM9515) and centrifuged. Pellet was rinsed in 70% ethanol, resuspended in Rnase-free water and samples were stored at -80°C.

For cDNA synthesis, we used the Super Script IV kit (*Thermo Fisher*, 18091200) according to the provided protocol. We adjusted the amount of RNA used to have the same for each replicate with a maximum amount of 2500 µg RNA. Samples were treated with RNase H to remove any trace of RNA for 20 min at 37°C and stored at - 20°C.

### Quantitative real-time PCR

Total RNA was extracted at 36 hpf from tails of control or pcdh18a i1-e2 MO injected embryos. cDNAs were synthetized and diluted at a final concentration of 15 µg/µl for each biological replicate. qPCR was performed using 7.5 µl of Takyon Rox SYBR Master mix blue dTTP kit (*Eurogentec*, UF-RSMT-B0701), 1.5 µl of cDNA and 0.5 µl of each 5µM primers (see Table 3). Real time qPCR was carried out for three biological replicates with measurements taken from three technical replicates on an Applied Biosystems 7300 Real Time PCR system (Thermofisher). After normalization with zebrafish elongation factor 1a (ef1a), the relative expression of *fn1a* and *fn1b* was determined by using the delta-delta-Ct method.

**Table 3.**
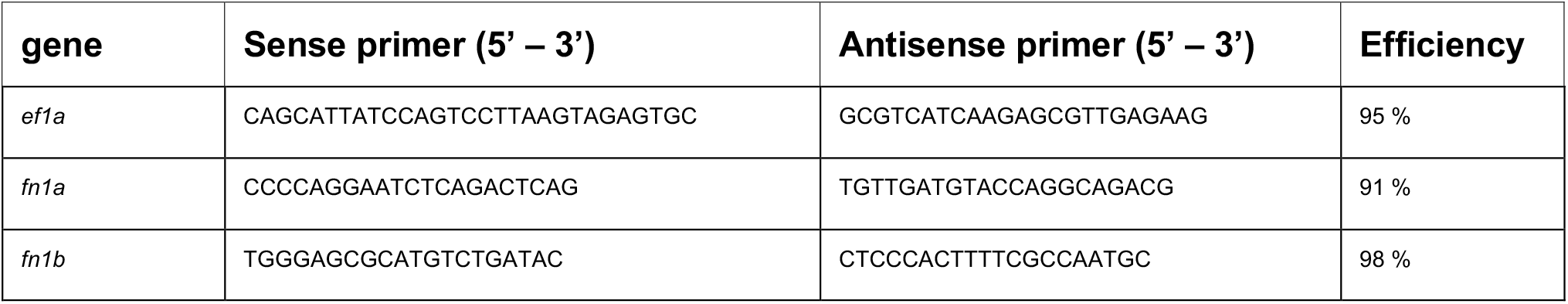
qPCR primers and efficiency measurement.

Primer efficiency measurement was done by diluting a reference batch of cDNA at 5x, 10x, 20x, 40x and performing qPCR as described previously. Calibration curves were obtained with the delta-delta-Ct method and the formula (10^(−1/slope)-1^)x100 to obtain the efficiency. Only primers with an efficiency comprised between 90 and 110%, with no amplification in H2O negative control, were selected.

### Assessment of pcdh18a morpholino specificity and efficiency

To verify that the pcdh18a i1e2 MO prevents splicing of the first intron of the *pcdh18a* gene, we performed PCR on cDNA from control or pcdh18a MO injected embryos. We used one primer in exon1 and one in exon2: FP 5’ – CAGTCGTCAATCCCTGAACAG 3’ and RP 5’ – GCTGCGAGAGTATTTGTTCCC – 3’, or one primer in exon1 and one in exon3: FP 5’–CAGTCGTCAATCCCTGAACAG–3’ and RP 5’– CGCCACGCCCACTATCTTTC–3’ and checked that the size of DNA fragments of morphants corresponded to the insertion of intron1 compared to controls. Sequencing of pcdh18a i1e2 morphant cDNA confirmed intron1 retention.

### Embryo mounting for confocal imaging

Dechorionated and anesthetized embryos were mounted in lateral position with their tails in close contact with the bottom of a µ-Dish 35 mm glass-bottom (*Ibidi*, 81158). They were embedded in 1x low-melting agarose (*Promega*, V2111) in 1x tricaine /1x PTU Volvic water, which was also added on top of it after agarose solidification.

### In vivo confocal imaging

For *in vivo* live imaging we used an inverted spinning disk confocal microscope (Andor, Resolution WD, Yokagawa CSU-W1, coupled with a Leica Dmi8 microscope), equipped with 488 nm and 561 nm laser diodes for excitation, 2 digital CMOS cameras (Hamamatsu ORCA-Flash4.0 LT, 2048×2048 pixels), a 63x water immersion objective (Leica HC PL APO CS2, NA 1.20, WD 300 μm) and the MetaMorph^®^ acquisition software. Optical planes were spaced by 0.3 µm and laser diode power was set at 10% with exposure time between 100 and 300 ms, depending on the transgenic line used. For *in vivo* time lapse imaging, we used an inverted laser scanning confocal microscope (Leica TCS SP8, CTR 6500), equipped with 488 nm and 552 nm lasers for excitation, 2 PMT and a Hybrid Detector, and a 40x or 20x immersion objective (Leica HC PL APO CS2 NA 1.1, WD 650 μm, and NA 0.75, WD 670 μm respectively) and the LAS X acquisition software. Usually, 2 controls and 2 morphants were acquired in parallel every 10 min with optical planes spaced by 1.5 µm.

For immunohistochemistry imaging, we used an inverted scanning confocal microscope (Leica TCS SPE, CTR 6000), equipped with 405 nm, 488 nm, 532 nm and 635 nm lasers for excitation, 1 PMT detector, a Leica ACS APO 40x oil objective (NA 1.15, WD 270 μm) and the LAS X acquisition software. Optical planes were spaced by 0.8 µm.

### VE-DIC microscopy

VE-DIC (for “video-enhanced differential interference contrast microscopy”) acquisitions of in situ hybridizations were performed using a Polyvar 2 microscope (Reichert) and a tri-CCD HV-D25 analog camera (Hitachi), adjusted to maximize contrast (Herbomel and Levraud, 2005). Embryos were imaged with either a 10x objective (Reichert PlanFIApo, NA 0.30), or a 40x water lens (Olympus), LUMPlanFi, NA: 0.80. Image sequences were recorded on Mini DV digital video tapes (Sony), using a GV-D1000E PAL recorder (Sony), then the selected images were extracted using the SwiftCapture software (Ben Software).

### Image analysis

For basic image treatment, we used the Fiji software (PMID 22743772) and for 3D visualization we used the Imaris software (*Bitplane*, v8.1). Cell tracking was made by using the manual tracking function of TrackMate plugin (v.6.0.3) of Fiji and plotted with a Python script written by S. Rigaud (Image Analysis Hub of Institut Pasteur). For filopodia analysis, we used the Filopodyan plugin (v1.2.6) available on Fiji (Urbancic et al., 2017). The cut-off threshold was set to 1 µm. For each condition, 10 SCP leader cells were analyzed during 10 time points, corresponding to 90 min of cell migration, from confocal time lapse sequences of Tg(*cspg4:Gal4; UAS:Lifeact-GFP*) embryos injected with control or pcdh18a i1-e2 MO. For Filopodyan setup, we used the triangle threshold, 2 or 3 ID iterations and from 1 to 3 LoG sigma, allowing to detect the angles of filopodia relative to the imaged field and their length for each cell. The direction of cell migration was derived from the initial and final position of the cell in the analyzed time window, then a Python script written by D. Ershov (Image Analysis Hub, Institut Pasteur) derived the angle of filopodia relative to the direction of cell migration from 0° to 180° and their length.

### Statistical analysis

Basic statistical analysis was performed using GraphPad Prism 9 software. To compare control to morphant groups, we first verified if the data followed a normal distribution by using the Shapiro-Wilk test. If it was the case, we used a Student t-test assuming that both populations have the same SD, if not we used the non-parametric Wilcoxon-Mann-Whitney test. For the homogeneity of the analyses, if one data set followed a normal distribution and the other not, in the same experimental series, we applied the non-parametric test for both.

Filopodyan analysis was performed using a R-script written by H. Julienne. Because the same cell is followed during 10 time points to get an average of the relative angle for each cell, results are not independent. A linear mixed effect model was implemented to add a random variable effect, allowing to model the effect of repeated measurements for each cell.

## Acknowledgements

We thank Stephane Rigaud, Dmitry Ershov and Jean-Yves Tinevez (Image Analysis Hub of Institut Pasteur) and Hannah Julienne (Bioinformatics & Biostatistics Hub of Institut Pasteur) for implementation of Python scripts and a R-script, respectively, that helped us quantify filopodia dynamics and statistics.

## References

Aamar, E. and Dawid, I. B. (2008). Protocadherin-18a has a role in cell adhesion, behavior and migration in zebrafish development. Dev Biol 318, 335–346.

Asakawa, K. and Kawakami, K. (2018). Protocadherin-Mediated Cell Repulsion Controls the Central Topography and Efferent Projections of the Abducens Nucleus. Cell Rep. 24, 1562–1572.

Biswas, S., Emond, M. R. and Jontes, J. D. (2010). Protocadherin-19 and N-cadherin interact to control cell movements during anterior neurulation. J. Cell Biol. 191, 1029–1041.

Biswas, S., Emond, M. R., Duy, P. Q., Hao, L. T., Beattie, C. E. and Jontes, J. D. (2014). Protocadherin-18b interacts with Nap1 to control motor axon growth and arborization in zebrafish. Mol. Biol. Cell 25, 633–642.

Bosze, B., Ono, Y., Mattes, B., Sinner, C., Gourain, V., Thumberger, T., Tlili, S., Wittbrodt, J., Saunders, T. E., Strähle, U., et al. (2020). Pcdh18a regulates endocytosis of E-cadherin during axial mesoderm development in zebrafish. Histochem. Cell Biol. 154, 463–480.

Chen, X., Koh, E., Yoder, M. and Gumbiner, B. M. (2009). A Protocadherin-Cadherin-FLRT3 Complex Controls Cell Adhesion and Morphogenesis. PLoS One 4, e8411.

Chen, B., Brinkmann, K., Chen, Z., Pak, C. W., Liao, Y., Shi, S., Henry, L., Grishin, N. V, Bogdan, S. and Rosen, M. K. (2014). The WAVE regulatory complex links diverse receptors to the actin cytoskeleton. Cell 156, 195–207.

Chi, N. C., Shaw, R. M., De Val, S., Kang, G., Jan, L. Y., Black, B. L. and Stainier, D. Y. R. (2008). Foxn4 directly regulates tbx2b expression and atrioventricular canal formation. Genes Dev. 22, 734–739.

Clark, A. G. and Vignjevic, D. M. (2015). Modes of cancer cell invasion and the role of the microenvironment. Curr. Opin. Cell Biol. 36, 13–22.

Gao, X., Xu, C., Asada, N. and Frenette, P. S. (2018). The hematopoietic stem cell niche: from embryo to adult. Development 145, dev139691.

Harrison, O. J., Brasch, J., Katsamba, P. S., Ahlsen, G., Noble, A. J., Dan, H., Sampogna, R. V., Potter, C. S., Carragher, B., Honig, B., et al. (2020). Family-wide Structural and Biophysical Analysis of Binding Interactions among Non-clustered δ-Protocadherins. Cell Rep. 30, 2655–2671.e7.

Hayashi, S., Inoue, Y., Kiyonari, H., Abe, T., Misaki, K., Moriguchi, H., Tanaka, Y. and Takeichi, M. (2014). Protocadherin-17 Mediates Collective Axon Extension by Recruiting Actin Regulator Complexes to Interaxonal Contacts. Dev. Cell 30, 673–687.

Helker, C. S. M., Schuermann, A., Karpanen, T., Zeuschner, D., Belting, H.-G., Affolter, M., Schulte-Merker, S. and Herzog, W. (2013). The zebrafish common cardinal veins develop by a novel mechanism: lumen ensheathment. Development 140, 2776–2786.

Herbomel, P. and Levraud, J. P. (2005). Imaging early macrophage differentiation, migration, and behaviors in live zebrafish embryos. Methods Mol. Med. 105, 199–214.

Homayouni, R., Rice, D. S. and Curran, T. (2001). Disabled-1 interacts with a novel developmentally regulated protocadherin. Biochem. Biophys. Res. Commun. 289, 539–547.

Jontes, J. D. (2016). The nonclustered protocadherins. In The Cadherin Superfamily: Key Regulators of Animal Development and Physiology, pp. 223–249.

Kimmel, C. B., Ballard, W. W., Kimmel, S. R., Ullmann, B. and Schilling, T. F. (1995). Stages of embryonic development of the zebrafish. Dev. Dyn. 203, 253–310.

Komoriya, A., Green, L. J., Mervic, M., Yamada, S. S., Yamada, K. M. and Humphries, M. J. (1991). The minimal essential sequence for a major cell type-specific adhesion site (CS1) within the alternatively spliced type III connecting segment domain of fibronectin is leucine-aspartic acid-valine. J. Biol. Chem. 266, 15075–15079.

Kraft, B., Berger, C. D., Wallkamm, V., Steinbeisser, H. and Wedlich, D. (2012). Wnt-11 and Fz7 reduce cell adhesion in convergent extension by sequestration of PAPC and C-cadherin. J. Cell Biol. 198, 695–709.

Lee, R. T. H., Knapik, E. W., Thiery, J. P. and Carney, T. J. (2013). An exclusively mesodermal origin of fin mesenchyme demonstrates that zebrafish trunk neural crest does not generate ectomesenchyme. Dev. 140, 2923–2932.

Morishita, H. and Yagi, T. (2007). Protocadherin family: diversity, structure, and function. Curr. Opin. Cell Biol. 19, 584–592.

Murayama, E., Kissa, K., Zapata, A., Mordelet, E., Briolat, V., Lin, H. F., Handin, R. I. and Herbomel, P. (2006). Tracing Hematopoietic Precursor Migration to Successive Hematopoietic Organs during Zebrafish Development. Immunity 25, 963–975.

Murayama, E., Sarris, M., Redd, M., Le Guyader, D., Vivier, C., Horsley, W., Trede, N. and Herbomel, P. (2015). NACA deficiency reveals the crucial role of somite-derived stromal cells in haematopoietic niche formation. Nat. Commun. 6, 8375.

Nakao, S., Platek, A., Hirano, S. and Takeichi, M. (2008). Contact-dependent promotion of cell migration by the OL-protocadherin-Nap1 interaction. J. Cell Biol. 182, 395–410.

Pancho, A., Aerts, T., Mitsogiannis, M. D. and Seuntjens, E. (2020). Protocadherins at the Crossroad of Signaling Pathways. Front. Mol. Neurosci. 13, 117.

Parinov, S., Kondrichin, I., Korzh, V. and Emelyanov, A. (2004). Tol2 transposon-mediated enhancer trap to identify developmentally regulated zebrafish genes in vivo. Dev. Dyn. 231, 449–459.

Redies, C., Vanhalst, K. and Van Roy, F. (2005). δ-Protocadherins: Unique structures and functions. Cell. Mol. Life Sci. 62, 2840–2852.

Seger, C., Hargrave, M., Wang, X., Chai, R. J., Elworthy, S. and Ingham, P. W. (2011). Analysis of Pax7 expressing myogenic cells in zebrafish muscle development, injury, and models of disease. Dev. Dyn. 240, 2440–2451.

Stickney, H. L., Barresi, M. J. F. and Devoto, S. H. (2000). Somite development in zebrafish. Dev. Dyn. 219, 287–303.

Suster, M. L., Abe, G., Schouw, A. and Kawakami, K. (2011). Transposon-mediated BAC transgenesis in zebrafish. Nat. Protoc. 6, 1998–2021.

Tai, K., Kubota, M., Shiono, K., Tokutsu, H. and Suzuki, S. T. (2010). Adhesion properties and retinofugal expression of chicken protocadherin-19. Brain Res. 1344, 13–24.

Thisse, C. and Thisse, B. (2008). High-resolution in situ hybridization to whole-mount zebrafish embryos. Nat. Protoc. 3, 59–69.

Urbancic, V., Butler, R., Richier, B., Peter, M., Mason, J., Livesey, F. J., Holt, C. E. and Gallop, J. L. (2017). Filopodyan: An open-source pipeline for the analysis of filopodia. J. Cell Biol. 216, 3405–3422.

Vanhalst, K., Kools, P., Staes, K., Van Roy, F. and Redies, C. (2005). δ-Protocadherins: A gene family expressed differentially in the mouse brain. Cell. Mol. Life Sci. 62, 1247–1259.

Wakayama, Y., Fukuhara, S., Ando, K., Matsuda, M. and Mochizuki, N. (2015). Cdc42 mediates Bmp - Induced sprouting angiogenesis through Fmnl3-driven assembly of endothelial filopodia in zebrafish. Dev. Cell 32, 109–122.

Wolf, K., Mazo, I., Leung, H., Engelke, K., Von Andrian, U. H., Deryugina, E. I., Strongin, A. Y., Bröcker, E. B. and Friedl, P. (2003). Compensation mechanism in tumor cell migration: Mesenchymal-amoeboid transition after blocking of pericellular proteolysis. J. Cell Biol. 160, 267–277.

Wu, J., Jiang, J., Chen, B., Wang, K., Tang, Y. and Liang, X. (2021). Plasticity of cancer cell invasion: Patterns and mechanisms. Transl. Oncol. 14, 100899.

Xue, Y., Liu, D., Cui, G., Ding, Y., Ai, D., Gao, S., Zhang, Y., Suo, S., Wang, X., Lv, P., et al. (2019). A 3D Atlas of Hematopoietic Stem and Progenitor Cell Expansion by Multi-dimensional RNA-Seq Analysis. Cell Rep. 27, 1567–1578.e5.

